# Re-investigation of classic T cell subsets and identification of novel cell subpopulations by single-cell RNA sequencing

**DOI:** 10.1101/2021.02.11.430754

**Authors:** Xuefei Wang, Xiangru Shen, Shan Chen, Hongyi Liu, Ni Hong, Xi Chen, Wenfei Jin

## Abstract

Classic T cell subsets are defined by a small set of cell surface markers, while single cell RNA sequencing (scRNA-seq) clusters cells using genome-wide gene expression profiles. The relationship between scRNA-seq Clustered-Populations (scCPops) and cell surface marker-defined classic T cell subsets remain unclear. Here, we interrogated 6 bead-enriched T cell subsets with 62,235 single cell transcriptomes and re-grouped them into 9 scCPops. Bead-enriched CD4 Naïve and CD8 Naïve were mainly clustered into their scCPop counterparts, while cells from the other T cell subsets were assigned to multiple scCPops including mucosal-associated invariant T cells and natural killer T cells. The multiple T cell subsets that form a single scCPop exhibited similar expression pattern, but not vice versa, indicating scCPops are much homogeneous cell populations with similar cell states. Interestingly, we discovered and named IFN^hi^ T, a new T cell subpopulation that highly expressed Interferon Signaling Associated Genes (ISAGs). We further enriched IFN^hi^ T by FACS sorting of BST2 for scRNA-seq analyses. IFN^hi^ T cluster disappeared on tSNE plot after removing ISAGs, while IFN^hi^ T cluster showed up by tSNE analyses of ISAGs alone, indicating ISAGs are the major contributor of IFN^hi^ T cluster. BST2+ T cells and BST2− T cells showing different efficiencies of T cell activation indicates high level of ISAGs may contribute to quick immune responses.

## Introduction

T cells, also know as T lymphocytes, are an essential component of adaptive immunity for protecting us against various pathogens. In fact, T cells not only play a key role in elimination of invasive pathogens and cancer cells, but also are important targets in the treatment of autoimmune diseases (*1–5*). The major T cells can be classified into CD4 T cells and CD8 T cells based on cell functions and cell surface markers. CD4 T cells, also known as T helper cells (Th), play a central role in adaptive immune response to pathogens, such as assisting B cells to produce antibodies, recruiting granulocytes to infected sites, and producing cytokines and chemokine to orchestrate immune response (*6*). Many distinct CD4 T cell subsets have been identified since the identification of Th1 by Mosmann *et al.* (*6, 7*). CD25+ CD4+ T cells, namely regulatory T cells (Treg), suppress self-reactive lymphocytes to maintain immunologic self-tolerance (*8, 9*). CD8 T cells, also known as cytotoxic T lymphocytes (CTLs), are important for immune defense against intracellular pathogens (*10, 11*). On the other hand, naïve T cells egress from thymus and are fairly quiescent, which could differentiate into distinct functional T cell subsets after they interact with cognate antigens, while memory T cells protect us against previously encountered pathogens (*3*). These T cell subsets could be enriched or sorted out by fluorescence activated cell sorting (FACS) or magnetic beads since each of them is defined by a set of specific cell surface markers. Then population-average assays were routinely used to characterize the biochemical and molecular features of each T cell subset.

Single cell RNA-sequencing (scRNA-seq), profiling gene expression at single cell resolution, has become an ideal approach for identifying cellular heterogeneity, searching new/rare cell types, pseudotime inference and inferring regulators underlying lineage changes (*5, 12–15*). Most studies focused on searching cellular heterogeneity and new cell types in well-defined T cell subsets due to the high statistical power of scRNA-seq to distinguish cellular heterogeneities (*16–21*). Only few studies compared classic T cell subsets with scRNA-seq clustered-populations (scCPops), thus our knowledge about the cellular relationship between classic T cell subsets and scCPops is limited. It remains unknown whether we could find the scCPop counterpart for each classic T cell subset and whether classic T cell subsets could well represent their scCPop counterparts. Whether more T cell subpopulations could be identified by re-clustering multiple classic T cell subsets?

In this study, we integrated scRNA-seq data of 6 bead-enriched T cell subsets for investigating the cellular relationships between classic T cell subsets and scCPops. We found bead-enriched CD4 Naïve, CD8 Naïve and CD4 memory cells were mainly clustered into their scCPop counterparts, while cells from the other T cell subsets were assigned to multiple scCPops. The multiple T cell subsets that form a single scCPop exhibited similar expression pattern, but not vice versa, indicating scCPops are much homogeneous cell populations. We unexpectedly identified a new T cell subpopulation and named it IFN^hi^ T that highly expresses Interferon Signaling Associated Genes (ISAGs). BST2 was selected as a representative of ISAGs and BST2^+^ T cell showed quick T cell activation upon anti-CD3/CD28 co-stimulation, indicating high level of ISAGs may contribute to quick immune responses.

## Results

### T cells form single continuous entity on tSNE plot

The scRNA-seq data of PBMCs from a healthy donor (donor A) were obtained from Zheng et al (*22*). The data were visualized by t-Distributed Stochastic Neighbor Embedding (t-SNE) and clustered into 7 distinct cell clusters following our previous studies (*5, 15*) (fig. S1A). We identified the cell type of each cluster based on cluster-specific expressed genes (fig. S1A, S1B). The frequencies of major identified cell types in PBMCs are consistent with the expectations: 75.48% T cells (*CD3D* and *CD3E*), 8.35% natural killer cells (*KLRF1* and *NCR3*), 7.46% B cells (*CD79A* and *MS4A1*), 6.24% CD14 monocytes (*CD14* and *LYZ*) and 2.47% CD16 monocytes (*CD16* and *LYZ*). Although these T cells were clustered into Naïve T cells (T_N_) (*CCR7* and *SELL*), CD4 memory T cells (CD4 T_M_) (*CD4* and *GZMK*) and CD8 memory T cells (CD8 T_M_) (*CD8A* and *GZMK*), those T cell subpopulations formed a single continuous entity without obvious boundaries (fig. S1A).

### Bead-enriched T cell subsets on tSNE plot

The scRNA-seq data of 6 bead-enriched T cell subsets, namely CD4+ T helper cells (CD4 Th; containing all sorts of CD4 T cells), CD4+/CD45RA+/CD25− naive T cells (CD4 naïve), CD4+/CD45RO+ memory T cells (CD4 memory), CD4+/CD25+ regulatory T cells (CD4 Treg), CD8+ cytotoxic T cells (CD8 CTL; containing all all sorts of CD8 T cells) and CD8+/CD45RA+ naive cytotoxic T cells (CD8 Naïve), were obtained from Zheng et al (*22*) (Table S1). Analyses of each T cell subset showed there was no obvious subpopulation in each T cell subset except CD8 CTL (fig. S1C), which essentially consist with original report that these T cell subsets are relative pure (*22*). The 62,235 cells from the 6 bead-enriched T cell subsets were projected on tSNE plot and formed a single connected entity (Fig. 1A). The CD4 T cell subsets, namely CD4 Naïve, CD4 memory, CD4 Treg and CD4 Th, are mixed together on tSNE plot. The 6 bead-enriched T cell subsets are overlapped with the T cells from PBMCs on tSNE plot (Fig. 1B), indicating the cellular heterogeneity among T cells are quite small comparing with that between T cells and other cell types. Highlighting the T cell subsets one by one showed that each T cell subset was concentrated on specific locations of the tSNE plot (Fig. 1C).

**Fig. 1.**
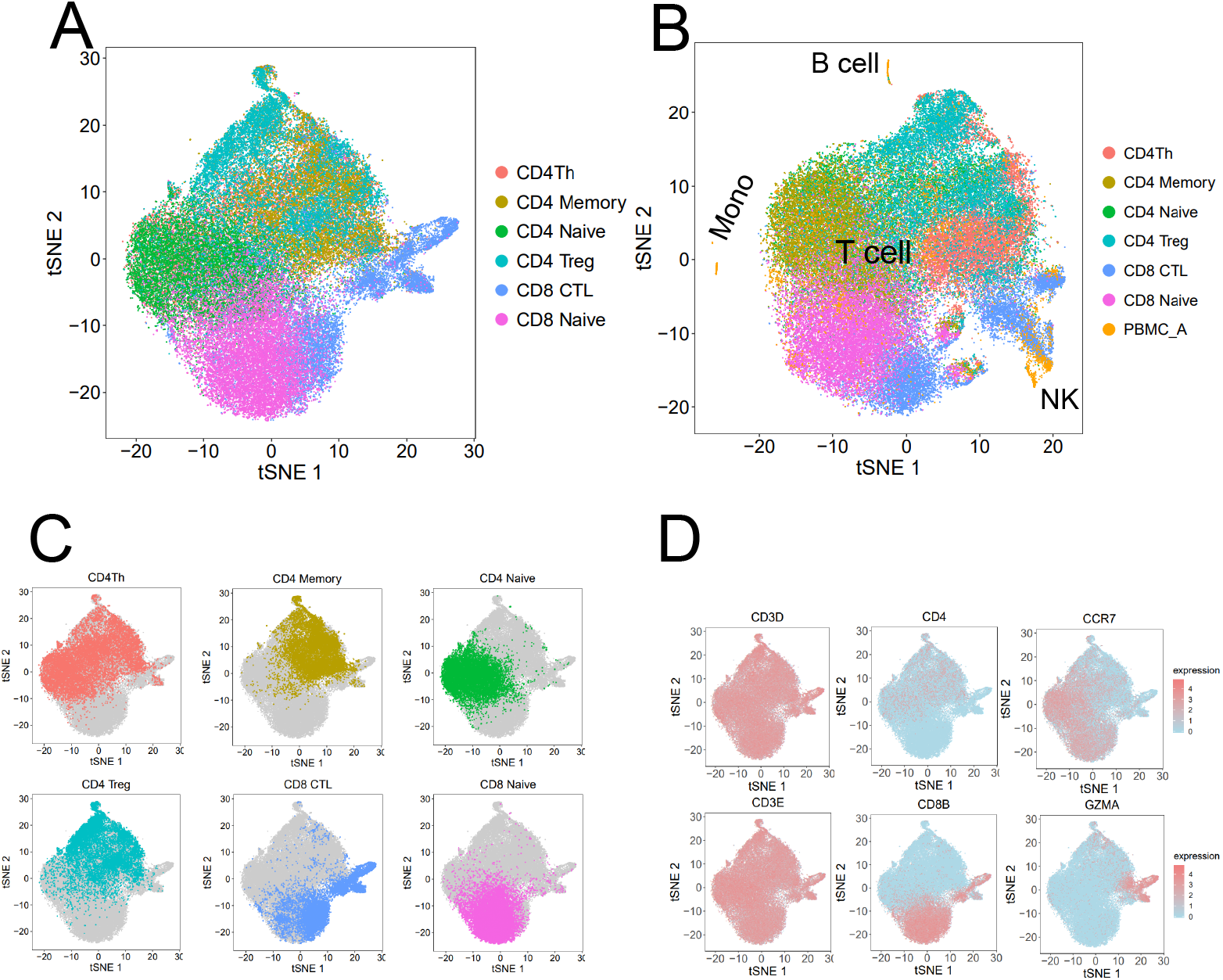
The 6 bead-enriched T cell subsets and their features. A. tSNE projection of the 6 bead-enriched T cell subsets, colored by each T cell subset. B. tSNE projection of PBMCs and the 6 T cell subsets, the PBMC and 6 T cell subsets colored differently. C. Distribution of T cell subset in tSNE plot of the 6 bead-enriched T cell subsets, highlighting one T cell subset in each panel. D. tSNE projection of the 6 T cell subsets, with each cell colored based on their normalized expression of *CD3D*, *CD3E*, *CD4*, *CD8B*, *CCR7* and *GZMA*, respectively.

*CD3D* and *CD3E*, the T cell marker genes, are expressed in each T cell subset (Fig. 1D). *CD4* and *CD8B* are highly expressed in the upper part and lower part of the tSNE plot, respectively (Fig. 1D), consistent with the observation that CD4 and CD8 T cell subsets are in upper part and lower part of tSNE plot, respectively. Expression of *CCR7* and *GZMA* indicate the Naïve/Naïve-like T cells are on the left of tSNE plot and cells secreting cytotoxic protein are on the right of tSNE plot, respectively. Overall, expression of marker genes on tSNE plot indicates that clustering cells by scRNA-seq data could provide further biological insights.

### scRNA-seq Clustered Populations (scCPops)

The 62,235 T cells from 6 the bead-enriched T cell subsets were clustered into 9 clusters/scCPops by Seurat(*23, 24*) analyses of the scRNA-seq data (Fig. 2A). The cell type of each scCPop was inferred based on scCPop-specific highly expressed genes (Fig. 2B, 2C, fig. S2A, S2B, Table S2). The scCPops were named after their classic T cell counterpart with ‘sc’ prefix. Among the 9 T cell scCPops, five of which exhibit strong signals matching well-known classic T cell subsets: CD4 Naïve T cell (scCD4 T_N_; *CD4*, *CCR7*, *LEF1*, *SELL*), CD8 Naïve T cell (scCD8 T_N_; *CD8B*, *CCR7*, *LEF1*, *SELL*), CD4 memory T cells (scCD4 T_M_; *CD4*, *TNFRSF4*, *TIMP1*, *USP10*), CD8 effector memory T cell with CD45RA or CD8 effector cells (scCD8 T_EMRA_/scCD8 T_EFF_; *CD8B*, *FGFBP2*, *GZMH*, *GZMB*) and mucosal-associated invariant T cell (scMAIT; *KLRB1*, *NCR3*, *ZBTB16*, *SLC4A10*, *RORC*). Two scCPops exhibit strong signals matching CD4 regulatory T cell (*FOXP3*, *IL2RA*, *CCR10*), which were designated as CD4 regulatory T cell #1 (scTreg#1) and CD4 regulatory T cell #2 (scTreg#2), respectively. Compared to scTreg#1, scTreg#2 are significantly higher expressed *MIR4435-1HG*, *LINC00152*, *KLRB1*, *LIMS1*, *NOSIP* and *CD59* (fig. S2C), indicating scTreg#2 is in active proliferation (*25, 26*). The high expression of *TIMP1* and *TNFRSF4* in scCD4 T_M_ potentially explains the long life of memory T cells since both genes have anti-apoptotic function(*27, 28*).

**Fig. 2.**
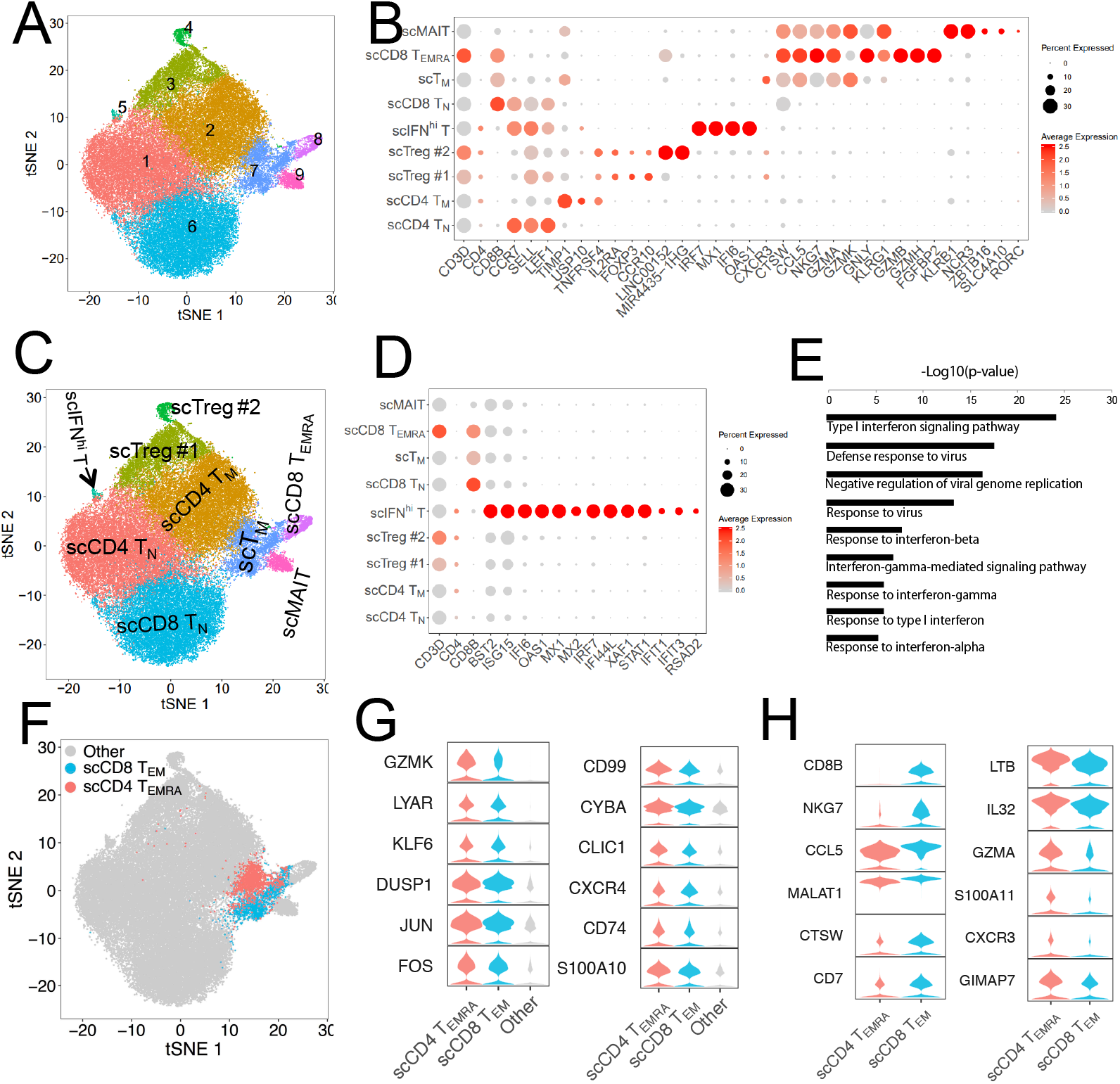
The 9 scRNA-seq Clustered Populations (scCPops) and their features. A. tSNE projection of the 62,235 T cells, where each cell is grouped into one of the 9 clusters distinguished by colors. Numbers at the panel indicate cluster identifier. B. Normalized expression level and expression percentage of the cell type specific genes in the 9 scCPops. C. tSNE projection of the 62,235 T cells, colored by the 9 scTCPops, with inferred cell types on the panel. D. Dotplot of scIFN^hi^ specific expressed genes. E. GO enrichment analysis of scIFN^hi^ specific expressed genes. F. The number of CD4 T cells and CD8 T cells almost equally contributed to scT_M_. G. scT_M_ specific expressed genes in its CD4 T subset and its CD8 T subset. H. Differentially expressed genes between CD4 T subset and CD8 T subset of scT_M_.

We identified 56 significantly high expressed genes in scCPop #5 by comparing scCPop #5 with the other cells (Table S2). These most significant scCPop #5 specific genes include *ISG15*, *IFI6*, *OAS1*, *MX1*, *MX2*, *IRF7*, *IFI44L*, *XAF1*, *STAT1*, *IFIT1*, *IFIT3* and *BST2* (Fig. 2B, 2D), which does not match any pre-defined T cell subset. Gene ontology (GO) analyses of the 56 genes showed the most significantly enriched GO terms were type I interferon signaling pathway (4.9×10^−25^), defense response to virus (1.8×10^−18^), negative regulation of viral genome replication (3.1×10^−17^), response to virus (3.5×10^−14^), response to interferon-beta (1.1×10^−8^), interferon-gamma-mediated signaling pathway (9.0×10^−8^) (Fig. 2E). We named this scCPop as interferon signaling high T cell (scIFN^hi^ T) since all those significantly enriched GO terms belong to interferon signalings. The discovery of scIFN^hi^ T in bead-enriched T cell subsets further pinpointed the cellular heterogeneity in well-defined classic T cell subsets.

Different from the other scCPop either belonging to CD4 T cell subset or CD8 T cell subset, scCPop #7 contains 41.39% CD4 T cells and 58.61% CD8 T cells. This scCPop was named scT_M_ because it highly expressed memory T cells specific genes such as *CXCR3, GZMK, TIMP1, LYAR*, *KLF6*, *DUSP1*, *JUN, FOS,* and *IL2RB* (Fig. 2B, 2G). The CD8 T cell in scT_M_ are highly expressed *NKG7*, *CCL5*, *CTSW*, *CD7* (Fig. 2H), indicating those T cells are CD8 effector memory T cell (scCD8 T_EM_); while CD4 T cells are highly expressed *IL32, LTB, GZMA, S100A11, CXCR3* and *GIMAP7* (Fig. 2H), indicating those cells are CD4 effector memory T cell re-expressing CD45RA (scCD4 T_EMRA_) or cytotoxic T lymphocytes (scCD4-CTL). It is well known that CD4 T cells and CD8 T cells diverged into two functional different lineages after double positive T cells (*4*). The scCD4 T_EMRA_/scCD4-CTL and scCD8 T_EM_ were clustered into one subpopulation (scT_M_) potentially because both cell subsets express cytotoxic associated genes. The observation that CD4 T cells and CD8 T cells could differentiate into similar phenotypes could be a model of convergence differentiation.

### Cellular relationships between classic T cell subsets and scCPops

The relationship between classic T cell subsets and scCPops was demonstrated by Sankey diagram (Fig. 3A). Bead-enriched CD4 Naïve and CD8 Naïve mainly clustered into their counterparts in scCPops, namely scCD4 T_N_ and scCD8 T_N,_ respectively, indicating the consistence between classic T cell enrichment and scRNA-seq clustering on those cell populations. These bead-enriched T cells solely clustering into their scCPops counterparts also indicated those T cell subsets are relatively homogeneous. On the other hand, CD4 Treg and CD4 memory were assigned to multiple scCPops, indicating high cell heterogeneity in the two classic T cell subsets. For instance, Treg was mainly assigned to 5 different scCPops, namely scCD4 T_N_, scCD4 T_M_, scT_M_, scTreg#1, and scTreg#2. As expected, CD4 Th and CD8 CTL contributed all CD4 scCPops and all CD8 scCPops, respectively. For instance, CD8 CTL was mainly clustered into scCD8 T_N_, scCD8 T_EMRA_, scT_M_ and scMAIT.

**Fig. 3.**
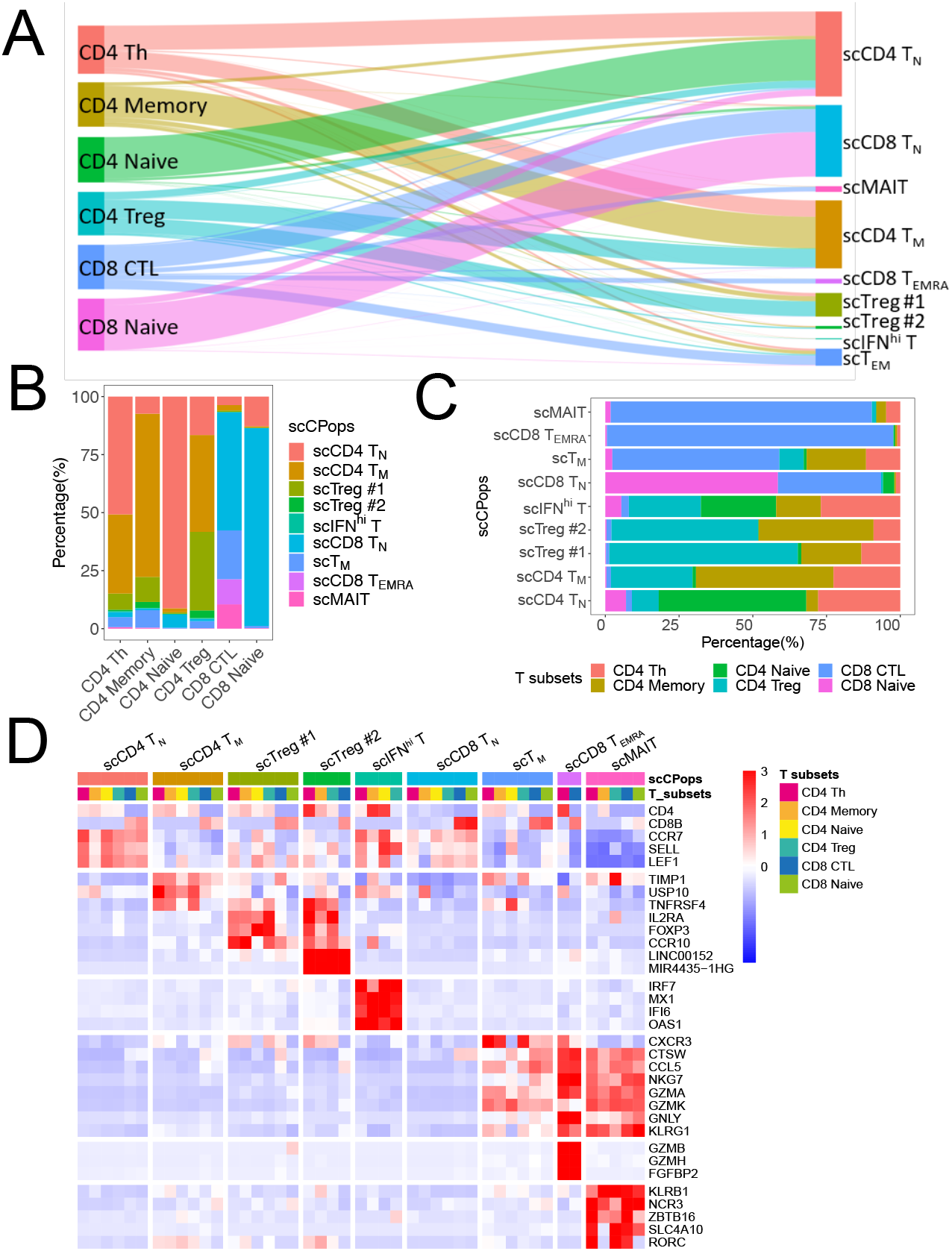
Relationships between bead-enriched T cell subsets and scCPops. A. Cellular relationships between the 6 bead-enriched T cell subsets and the 9 scCPops by Sankey diagram. Each node represents a cell populations and width of line between two nodes represents number of shared cells. B. Cell compositions of bead-enriched T cell subsets by bar chart. C. Bar plot displays cell compositions of scCPops based on bead-enriched T cell subsets. D. Gene expression heatmap of cell subpopulations generated by combinations of scCPops and bead-enriched T cell subsets.

Cell composition analyses of bead-enriched T cell subsets based on scCPops directly revealed the cellular heterogeneity in classic T cell subsets. CD4 Naïve consists of 91.21% scCD4 T_N_, 5.56% scCD8 T_N_, 1.74% scCD4 T_M_ and 0.63% scTreg#1 (Fig. 3B). CD8 Naïve consists of 85.23% scCD8 T_N_ and 12.79% scCD4 T_N_. CD4 memory consists of 70.35% scCD4 T_M_, 10.79% scTreg#1, 7.47% scT_M_ and 7.35% scCD4 T_N_ (Fig. 3B). CD4 Treg consists of 41.72% scCD4 T_M_, 33.86% scTreg#1, 16.57% scCD4 T_N_, 3.26% scTreg#2 and 3.07% scT_M_ (Fig. 3B). CD4 Th consists of 50.75% scCD4 T_N_, 34.11% scCD4 T_M_, 6.98% scT_Reg_#1, 4.32% scT_M_, and 2.14% scCD8 T_N_ (Fig. 3B). CD8 CTL consists of 50.96% scCD8 T_N_, 21.03% scT_M_, 10.82% scCD8 T_EMRA_, 10.49% scMAIT, 3.66% scCD4 T_N_ and 2.31% scCD4 T_M_, which should be the most heterogeneous among the 6 bead-enriched T cell subsets.

We further calculated the cell composition of scCPops based on classic T cell subsets. Both scMAIT and scCD8 T_EMRA_ were almost solely derived from CD8 CTL (Fig. 3A, 3C). However, the cell compositions of the other 7 scCPops were quite complex, with each scCPop deriving from several bead-enriched T cell subsets (Fig. 3C). In particular, all the 6 bead-enriched T cell subsets contributed to scIFN^hi^ T, arranging from 2.54% to 26.92% (Fig. 3C). These results indicated scIFN^hi^ T was a rare cell population accounting 0.26% of the analyzed cells, with CD4 T cell subsets containing high fraction of scIFN^hi^ T. scIFN^hi^ T accounts for about 0.4% of CD4 Naïve, CD4 Th and CD4 Treg, and 0.24% of CD4 memory, whereas only accounts for 0.04% CD8 CTL and 0.09% CD8 Naïve (fig. S2D).

The T cells were separated into 54 cell subpopulations by combinations of 9 scCPops and 6 bead-enriched T cell subsets. Finally, 45 cell subpopulations were left after we removed the subpopulations with number of cells <10. Cell subpopulations deriving from different bead-enriched T cell subsets while belonging to same scCPop always showed similar expression pattern (Fig. 3D). On the other hand, cell subpopulations deriving from different scCPops while belonging to same bead-enriched T cell subset usually showed quite different expression pattern (Fig. 3D, fig. S2E). These results are consistent with our pre-assumption that scCPops better represent the cell states than that of bead-enriched T cell subsets.

### Fine analyses of CD4 T cell subsets

The CD4 T cell subsets (fig. S3A) were further analyzed to better understand the subpopulations and lineages of CD4 T cells. The 40,696 cells from CD4 Th, CD4 naïve, CD4 memory and CD4 Treg formed a single connected entity on tSNE plot and different CD4 T cell subsets mixed together (Fig. 4A). We clustered these T cells into 6 scCPops (Fig. 4B, 4C), among which 4 scCPops have been identified aforementioned, namely scCD4 T_N_ (*CCR7*, *SELL*, *LEF1*), scIFN^hi^ T (*OAS1*, *IFI6*, *MX1*, *BST2*), scCD4 T_EMRA_ (*GZMA*, *CCL5*, *GZMK*, *CST7*, *LYAR*) and scTreg (*FOXP3*, *IL2RA*, *IL10RA*, *CD59*). The original scCD4 T_M_ were separated into central memory T cell (scCD4 T_CM_) (*CCR7*, *SELL*, *TCF7*) and effector memory T cell (scT_EM_) (*CCR7^−^*, *TIMP1*, *LGALS1*, *USP10*), respectively (Fig. 4B, 4C). Sankey diagram showed the cellular relationships between the 4 bead-enriched CD4 T cell subsets and the 6 scCPops were consistent with aforementioned analyses of all T cells. Briefly, bead-enriched CD4 Naïve almost solely clustered into its counterpart scCD4 T_N_. On the other hand, CD4 Th, CD4 memory and CD4 Treg contributed to all the 6 scCPops (Fig. 4D).

**Fig. 4.**
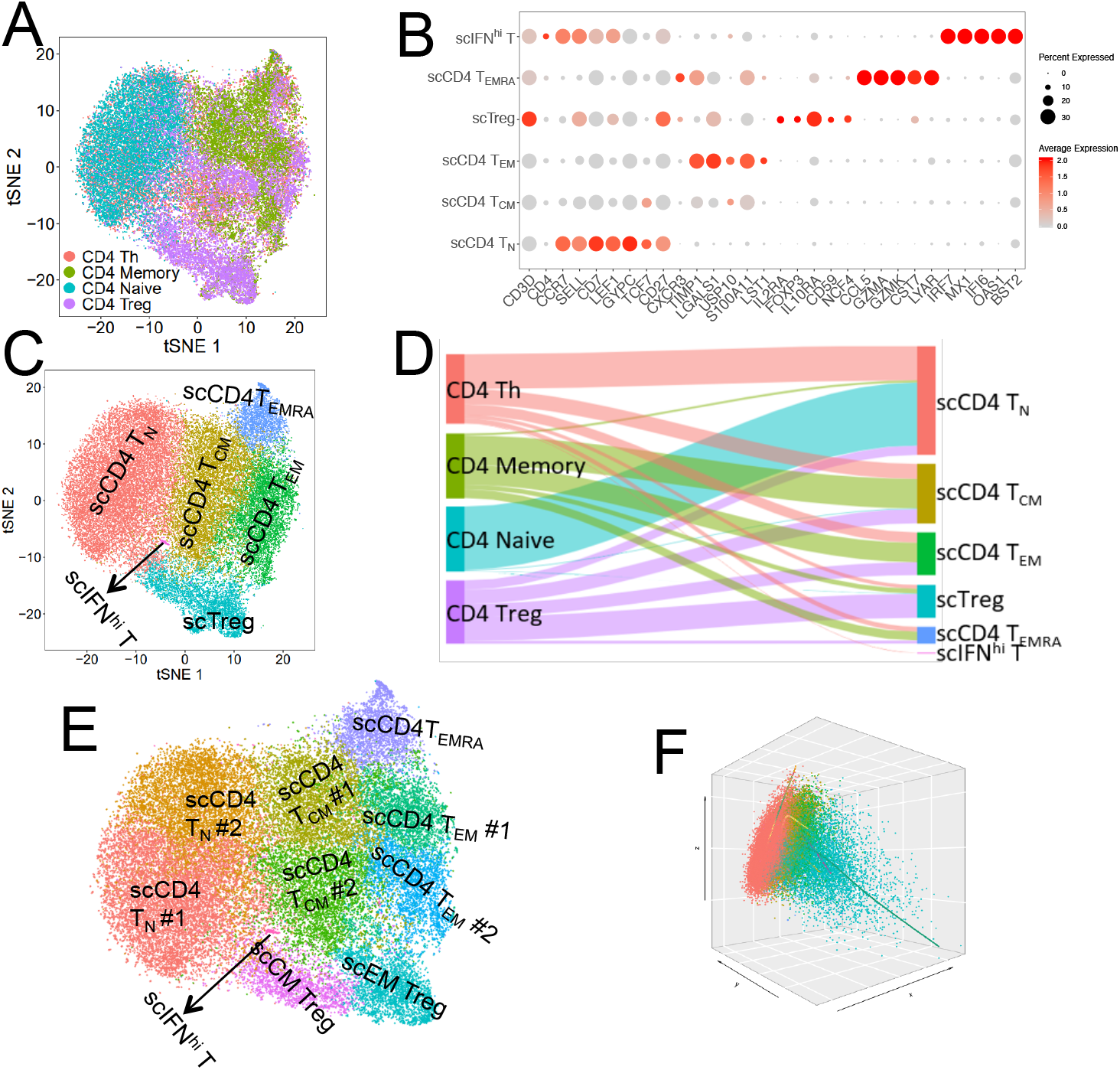
Reclassification of CD4 T cells and single cell trajectory. A. tSNE projection of CD4 T cells, with each cell colored according to bead-enriched T cell subsets that distinguished by colors. B. Normalized expression level and expression percentage of the cell type specific genes in 6 CD4 T cell scCPops. C. tSNE projection of CD4 T cells, where each cell is grouped into one of the 6 scCPops, with inferred cell types on the panel. D. Cellular relationships between the 4 CD4 T cell subsets and the 6 scCPops by Sankey diagram. Each node represents a cell populations and width of line between two nodes represents number of shared cells. E. The original 6 scCPops were further separated into 10 scCPops by setting high resolution. F. Single cell trajectories of CD4 T cells inferred by slingshot. The color of each scCPop is the same as that in Fig. 4C.

We further clustered the CD4 T cells into 10 scCPops for better understanding its fine substructure (Fig. 4E). The original scTreg was separated into central memory Treg (scCM Treg; *FOXP3*, *CCR7*, *LEF1*, *CTLA4^low^*) and effector memory Treg (scEM Treg; *FOXP3*, *IL2RA*, *LGALS1*, ANXA2, *CTLA4*) (fig. S3B). The original scCD4 T_N_ were separated into scCD4 T_N_#1 and scCD4 T_N_#2, with most CD4 naïve specific genes such as *CCR7*, *SELL*, *LEF1* and *TCF7* expressing higher in scCD4 T_N_#1 (fig. S3B). The original scCD4 T_EM_ were divided into scCD4 T_EM_#1 and scCD4 T_EM_#2 (Fig. 4E), with *KLRB1* and *PTGER2* expressing higher in scCD4 T_EM_#1, potentially indicating scCD4 T_EM_#1 is an intermediate state between scCD4 T_EM_#2 and scCD4 T_EMRA_. Sankey diagram showed that CD4 T, CD4 memory and CD4 Treg contributed to almost all the 10 fine scCPops, indicating the high cellar heterogeneity of bead bead-enriched T cells subsets (fig. S3C).

Application of Slingshot (*29*) on these CD4 T cells identifying 4 pseudotime lineages: 1) scCD4 T_N_ → scCD4 T_CM_ → scCD4 T_EM_ → scCD4 T_EMRA_; 2) scCD4 T_N_ → scCD4 T_CM_ → scCD4 T_EM_; 3) scCD4 T_N_ → scCD4 T_CM_ → scCD4 Treg; 4) scCD4 T_N_ → scIFN^hi^ (Fig. 4F). The CD4 T cell differentiations are a continuously process, in which Naïve cells first progressed into central memory-like cells from which it developed into different functional CD4 T cells. The first inferred lineage was consisted with recent reports on CD4 T cell differentiation(*20, 30, 31*), which is the most important lineage since it includes most of the CD4 T cells.

### Fine analyses of CTLs

We integrated 3 T cell populations, namely bead-enriched CD8 CTL, bead-enriched CD8 Naïve and scCD4 T_EMRA_/scCD4 CTLs that clustered into scT_M_ with CD8 T cells (fig. S4A), for detecting fine substructure in these T cell subsets and for providing insight into the relationship between CD4 CTLs and CD8 CTLs. The 3 CTL populations formed one major group and several small groups on tSNE plot, with the CD8 Naïve on the major group and scCD4 T_EMRA_ overlapping with some CD8 CTLs (Fig. 5A). We further clustered these T cells into 6 scCPops (Fig. 5B, 5C), namely scCD8 T_N_ (*CCR7*, *SELL*, *LEF*, *CD69^low^*), scCD8 T_CM_ (*CCR7*, *SELL*, *LEF1*, *CD69*), scT_M_ (*CXCR3*, *TRADD*, *GZMA*, *GZMK*, *CCL5*), scCD8 T_EMRA_ (*FGFBP2*, *GZMH*, *GZMB*, *GNLY*), scMAIT (*KLRB1*, *ZBTB16*, *NCR3*, *SLC4A10*, *RORC*) and Natural killer cell (scNKT; *NCR3*, *KLRC2*, *ZNF683*, *XCL1*) (Fig. 5B, 5C). The expression pattern and cell composition of scT_M_ is the same as the aforementioned scT_M_ in Fig. 2. The expression of commonly used T cell markers on tSNE plot showed obvious scCPops specific features (Fig. 5D). Volcano plot showed that both MAIT and NKT expressed a lot cell type specific genes (fig. S4B, S4C), further indicating that MAITs and NKTs are outliers in CD8 CTL.

**Fig. 5.**
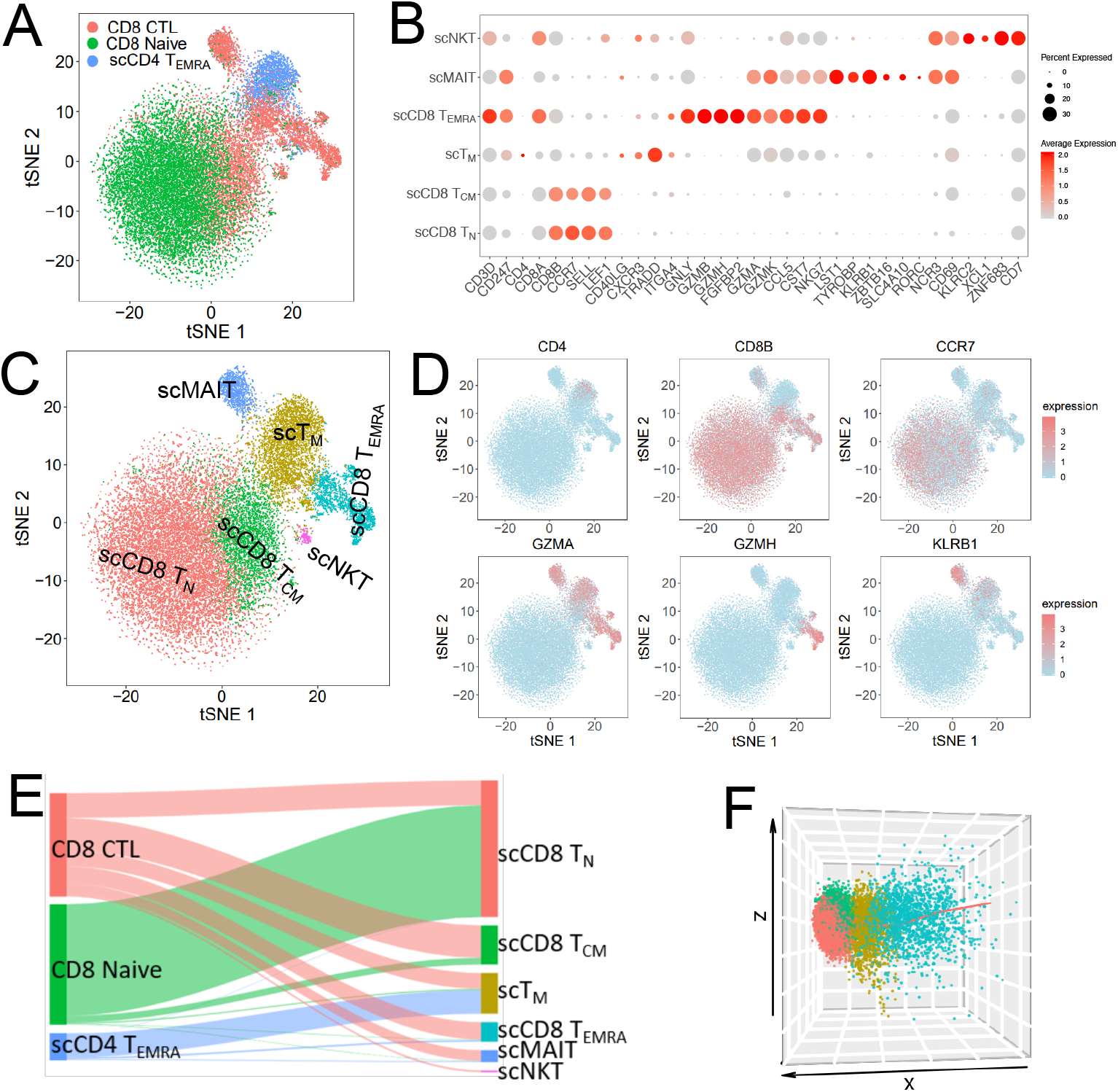
Reclassification of potential CTLs based on scRNA-seq and single cell trajectory. A. tSNE projection of CD8 Naïve, CD8 CTL and scCD4 T_EMRA_, with each cell colored according to cell populations that distinguished by colors. B. Normalized expression level and expression percentage of the cell type specific genes in the 6 scCPops. C. tSNE projection of CTLs, where each cell is grouped into one of the 6 scCPops, with inferred cell types on the panel. D. tSNE projection of CTLs, with each cell colored based on their normalized expression of *CD4*, *CD8B*, *CCR7*, *GZMA*, *GZMH* and *KLRB1*, respectively. E. Relationships between the initial T cell subsets and the 6 scCPops by Sankey diagram. Each node represents a cell populations and width of line between two nodes represents number of shared cells. F. Single cell trajectory of CD8 CTL inferred by slingshot. The color of each scCPops is the same as that in Fig. 5C.

Bead-enriched CD8 Naïve was mainly clustered into its counterpart scCD8 T_N_, with a small fraction clustered into scCD8 T_CM_ (Fig. 5E). scCD4 T_EMRA_, together with a fraction of cells from CD8 CTL, forms scT_M_, consistent with our aforementioned results. The cells from CD8 CTL contributed to each of the 6 scCPops (Fig. 5E), indicating multiple T cell subsets in commonly used CD8 CTL. Especially, the existence of un-ignorable amount of MAITs and NKTs in CD8 CTL indicated that the composition of classic CD8 CTL is complex.

We removed scMAIT, scCD4 T_EMRA_ and scNKT for inferring CD8 T cell differentiation lineage since the 3 cell populations do not belong to well-defined CD8 T cells. By conducting Slingshot (*29*), we inferred the pseudotime trajectory of CD8 T cell differentiation: 1) scCD8 T_N_ → scCD8 T_CM_ → scCD8 T_EM_ → scCD8 T_EMRA_ (Fig. 5F), which is similar to the CD4 main lineage. Based on inferred lineage and cell projection on PCA plot, the differentiation from CD8 Naïve T to scCD8 T_EMRA_ is a continuous process (Fig. 5F).

### Enrichment of IFN^hi^ T and identification of its features

Although *BST2* only showed weak scIFN^hi^ specificity (Fig. 2D, 2E), it is the only cell surface gene with well-established antibody among these ISAGs. We enriched scIFN^hi^ T by FACS sorting of BST2^hi^ cell from PBMC and made two independent scRNA-seq libraries (Fig. 6F, fig. S5A). These FACS sorted BST2 ^hi^ T cells was clustered into IFN^hi^ T, CD4 T_N_, CD8 T_N_, CD4 T_M_, CD8 T_M_, CD4 T_P_, MAIT, γδ T, NKT, eTreg and nTreg (Fig. 6B, fig. S5B). Although the enrichment of IFN^hi^ T has significantly increased, IFN^hi^ T only accounts for a moderate fraction of the BST2^hi^ cell because BST2 is expressed in many T cell subsets. The expression level of *BST2* in IFN^hi^ T is significantly higher than that of other cells based on the scRNAseq data (Fig. 6C). We further separated the non-IFN^hi^ T cells into *BST2^+^* T and *BST2^−^* T cells based on whether the expression of *BST2* was detected in scRNAseq data (Fig. 6D). The expression profile of IFN^hi^ T is quite different from that of *BST2^+^* T cells and *BST2^−^* T cells, while the expression profiles of *BST2^+^* T cells and *BST2^−^* T cells are more similar (Fig. 6E, 6F). IFN^hi^ T has 89 genes with significant higher expression, including *IF144L*, *MX1*, *IF1616*, *XAF1*, *ISG15*, *OAS3*, *IFI44*, *IRF7*, *MX2*, *IFITM1*, *OASL* and *IFI35*, than *BST2^+^* T cells (Fig. 6G). These IFN^hi^ T specific genes are significantly enriched GO terms including interferon signaling, defense response to virus, response to interferon γ and antiviral mechanism by IFN-stimulated genes (fig. S5C). In short, high expression of interferon signaling associated genes (ISAGs) is the most significant features of IFN^hi^ T since it significantly higher expressed ISAGs than *BST2^+^* cells.

**Fig. 6.**
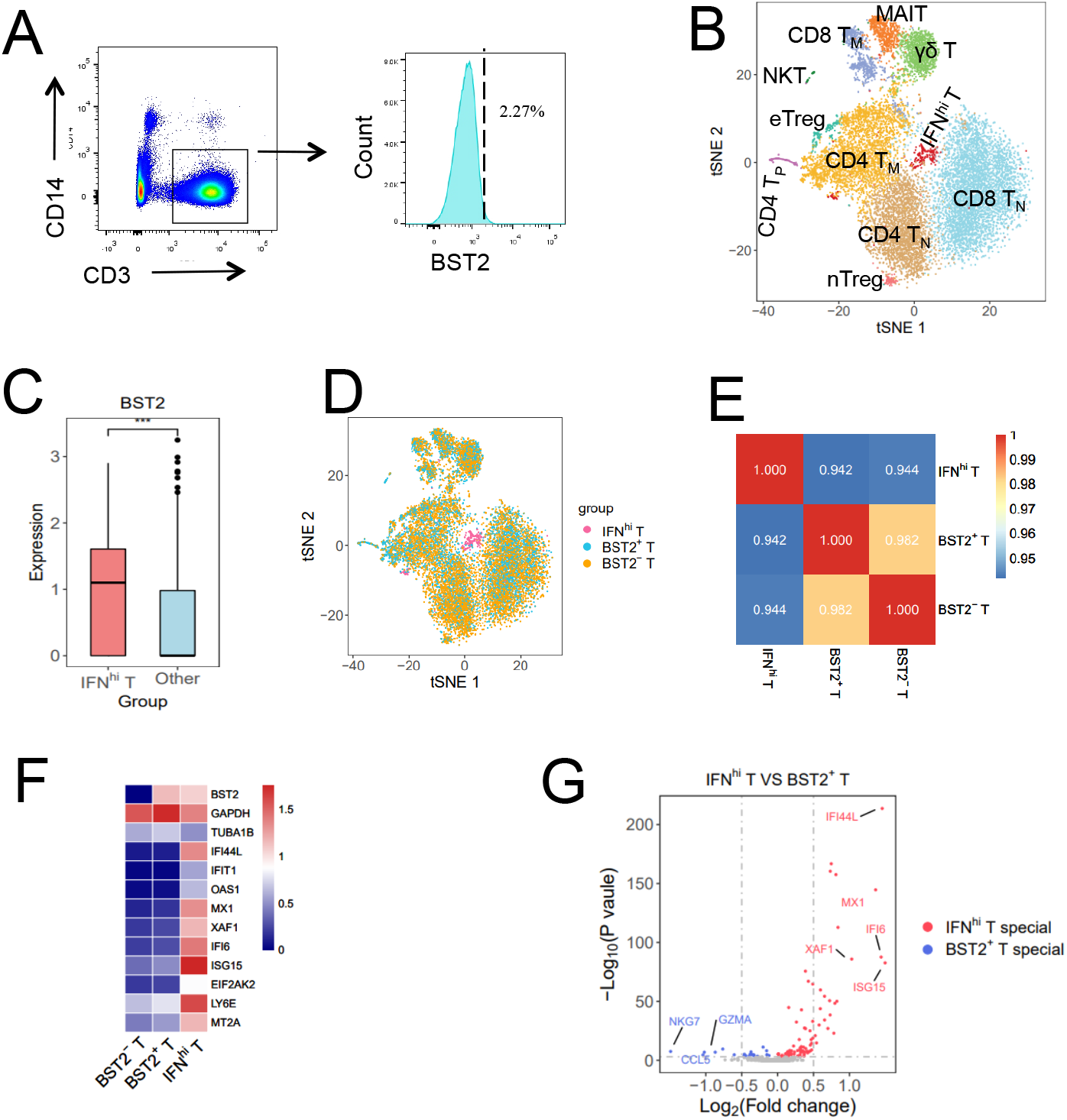
Enrichment of IFN^hi^ T by FACS sorting and feature of IFN^hi^ T. A. Sorting of BST2^hi^ T cells from PBMCs by FACS. B. tSNE projection of 13,644 sorted BST2^hi^ T cells, with each cell colored according to cell subpopulations. C. Box plot of *BST2* expression of IFN^hi^ T cells and other cells in BST2^hi^ T cells. *** represents a significant difference (P value <0.01). D. tSNE projection of BST2^hi^ T cells, with each cell colored according to IFN^hi^ T, BST2^+^ T and BST2^−^ T cells. *BST2^+^* T and *BST2^−^* T cells are defined by expression of BST2 in scRNA data. E. Spearman correlation coefficient between IFN^hi^ T, *BST2^+^* T and *BST2^−^* T. Color represents correlation coefficient between two groups. F. Heatmap of differentially expressed genes between IFN^hi^ T, *BST2^+^* T and *BST2^−^* T cells. Color represents average expression of genes in each group. G. Volcano plot for differentially expressed genes between IFN^hi^ T cell and *BST2^+^* T cell (P value <0.01). Red points represent IFN^hi^ T specific high expressed genes, while blue points represent *BST2^+^* T cell specific high expressed genes.

### ISAGs shape IFN^hi^ T and contribute quick T cell activation

We re-conducted tSNE analysis on BST2^hi^ T cells using gene set excluded ISAGs, the scCPops did not change except IFN^hi^ T disappeared (Fig. 7A). On the tSNE plot based on gene sets without ISAGs, IFN^hi^ T re-dispersed into the other scCPops, in particular concentrated on CD8 T_N_ (39.41%), CD4 T_N_ (28.39%) and CD4 T_M_ (28.39%); while only a few IFN^hi^ T dispersed into innate-like T cells such as MAIT, γδ T and NKT (Fig. 7B, fig. S5D). These results are expected since the expression pattern of IFN^hi^ T is much similar to CD4 T_N_, CD8 T_N_, and CD4 T_M_ (fig. S5B). Normalizations of the cell number of IFN^hi^ T to the cell number of its projecting scCPop showed CD8 T_N_, CD4 T_N_ and CD4 T_M_ had similar percentage of cells similar to IFN^hi^ T (fig. S5E). These results are consistent with our aforementioned observations that bead-enriched CD4 T cell subsets have much high fraction of cells that were clustered into IFN^hi^ T (fig. S2D). Interesting, IFN^hi^ T cluster showed up when we only used ISAGs to conduct tSNE analysis on BST2^hi^ T cells (Fig. 7C), further indicating that ISAGs are major contributor for the formation of IFN^hi^ T cluster.

**Fig. 7.**
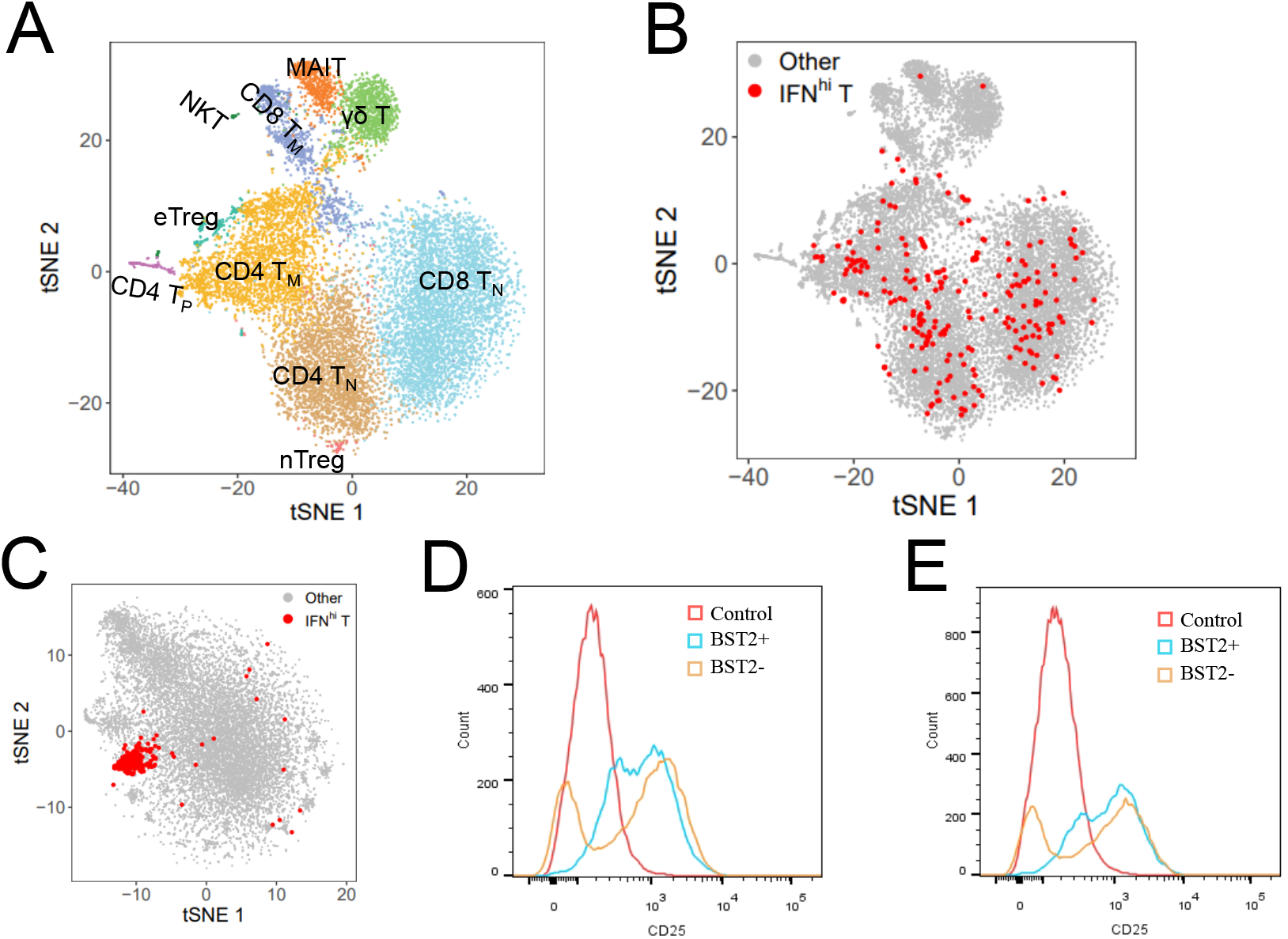
Interferon pathway associated genes (ISAGs) are the major contributor of IFN^hi^ T cell cluster and high level of ISAGs may contribute quick T cell activation. A. tSNE projection of BST2^hi^ T cells using gene sets that removed ISAGs, in which IFN^hi^ T cell cluster disappears. B. tSNE projection of BST2^hi^ T cells using gene sets that removed ISAGs, with red points highlighting the original identified IFN^hi^ T. C. tSNE projection of BST2^hi^ T cells using ISAGs, with red points highlighting original identified IFN^hi^ T. D-E. FACS analysis of CD25 showed almost all BST2+ T cells activated upon anti-CD3/CD28 beads for 12 hours, while only a fraction of BST2− T cells activated at the same time.

Constitutive expression of ISAGs plays pivotal roles in host responses to malignant cells and pathogens such as virus (*32–34*). We used FACS to sort the T cells into BST2+ T cells and BST2− T cells because BST2 level indicate activation level of ISAGs in some way. The sorted BST2+ T cells and BST2− T cells were activated by dynabeads^®^ human T-activator CD3/CD28 for 12 hours. FACS analysis of CD25 showed almost all BST2+ T cells become activation upon anti-CD3/CD28 co-stimulation, while only a fraction of BST2− T cells become activated (Fig. 7D, 7E, fig. S6). These results may indicate that almost all BST2+ T cells have being well prepared for T cell activation while only a fraction of BST2− T cells has being prepared for quick T cell activation.

## Discussion

It is assumed that each T cell subset has specific functions and several T cell subsets may coordinate with each other to complete a specific function (*35, 36*). The recent advance of scRNA-seq provided an unbiased approach to identify the T cell subpopulations and cell heterogeneity. Here we re-clustered the 6 bead-enriched T cell subsets using public scRNA-seq data. Among the 6 T cell subsets, CD4 Th and CD8 CTL include all sorts of CD4 T cells and all sorts of CD8 T cells, respectively, thus T cell subsets are well representative. However, Naïve T cells and memory T cells account majority of the T cells in this analyses (Fig 2B, 2C, 3A), and the signals from memory T cells and naïve T cells were strong. Therefore, we mainly focused on memory T cells and naïve T cells, while ignored other T cell subsets such as Th1, Th2 and Th17. We not only identified the scCPops corresponding to their bead-enriched counterparts, but also identified unexpected T cell subpopulations such as IFN^hi^ T, MAIT and NKT. Furthermore, we increased the resolution and identified much specific T cell subpopulation such as scCD4 T_CM_, scCD4 T_EM_, scCD4 T_EMRA_, scCM Treg, scEM Treg, scCD8 T_CM_, scCD8 T_EM_ and scCD8 T_EMRA_. These results showed that scRNA-seq clustering are essentially consistent with the classic FACS/bead-enriched approach. However, scRNA-seq clustering even identified rare cell subpopulation such as IFN^hi^ T and inferred much specific subpopulations, indicating its advantage comparing with the classic approach. Further analyses revealed that IFN^hi^ T showed quite unique features and may contribute to the quick T cell activation.

Although CD4 T cells and CD8 T cells essentially are distinguishable based on cell surface markers, the boundaries between CD4 T cells and CD8 T cells were blurred based on scRNA-seq data (Fig. 1A), indicating the overall transcriptional profiles between CD4 T cells and CD8 T cells were not completely different. In particular, we identified the scT_M_ cluster that contains both CD4 T cells and CD8 T cells. Furthermore, we inferred the trajectories of CD4 T cells and CD8 T cells, which showed that CD4 Naïve T cells and CD8 Naïve T cells differentiated into CD4 T_EMRA_ and CD8 T_EMRA_, respectively. Re-investigation of T cell subsets by scRAN-seq is of great significance for understanding T cell subsets, and helps us to further understand the mechanism of the human adaptive immune response, which also has important implications of the ongoing immunotherapy.

## Methods

### Isolation of peripheral blood mononuclear cells (PBMCs)

This study was approved by institutional review boards (IRBs) at Southern University of Science and Technology (SUSTech). All experiments were conducted following the protocols approved by IRBs at SUSTech. Human peripheral blood samples were obtained from a healthy adult blood donor. PBMCs were isolated by density-gradient sedimentation using Lymphoprep™ (Axis-Shield, Oslo, Norway) following the manufacture’s instructions. Briefly, peripheral blood sample was diluted with an equal volume of PBS containing 2% FBS, transferred into a Lymphoprep™ tube. Following centrifuged at 800 × g for 20 min at room temperature, the middle white phase was transferred to a fresh tube. The enriched mononuclear cells were washed with PBS containing 2% FBS and centrifuged at 500 × g for 5 min twice. Cell count and viability was assessed by Hemocytometer Counting.

### Isolation of T cells and T cell subpopulations

PBMCs were re-suspended in FACS buffer (PBS containing 2% FBS), and incubated with Human TruStain FcX™ (Fc Receptor Blocking Solution) (Biolegend, Cat#422302, USA) for 15 minutes on ice. After anti-human CD14-APC (61D3, eBioscience, Cat#17-0149-42, USA) and anti-human CD3-FITC (HIT3a, eBioscience, Cat#11-0039-42, USA) were added into PBMCs, they were incubated on ice for 30 minutes. These cells were washed twice with FACS buffer, and counter-stained with 1μg /mL DAPI. These cells were sorted on FACS Aria-III (Becton Dickinson) with CD14-CD3+ gating, thus we obtained the T cells. FlowJo (version 10) was used for FACS data analyses and Fig. plot. On the other hand, we simultaneously added anti-human CD317-PE (RS38E, BioLegend, Cat#348405, USA) during antibodies incubation for sorting BST2+ T cell or BST2^hi^ T cells.

### T cell activation

Following the manufacturer’s instructions, FACS sorted BST2+ T cells and BST2− T cells were stimulated with Dynabeads™ Human T-Activator CD3/CD28 for T Cell Expansion and Activation (ThermoFisher, cat#11161D, USA). Briefly, 8 × 10^4^ FACS sorted BST2+ T cells or BST2− T cells were suspended with 100–200 μL medium, respectively. Pre-washed Dynabeads^®^ was added at the bead-to-cell ratio of 1:1 and incubated in a humidified CO_2_ incubator at 37°C. After 12 hours, activated T cells were harvested by removal of Dynabeads^®^, and then stained with anti-CD25 (Cat #4302545, eBioscience) for FACS analyses of the efficiency of T cell activation.

### scRNA-seq library preparation and sequencing

FACS-sorted BST2+ T cells were directly processed for scRNA-seq with 10X Genomics 3′ kit v3 (10x Genomics, Pleasanton, CA) following manufacturer’s instructions. In short, about 1.6 × 10^4^ BST2+ T cells were loaded into single inlet of a 10x Genomics Chromium controller to generate the Gel Beads-in-Emulsions (GEMs). Two independent biological replicates were prepared, with each replicate recovering about 8000 cells. GEMs were used to complete the reverse transcription, cDNA amplification and library preparation. scRNA-seq libraries were prepared using Chromium Single Cell 3′ Library Construction Kit v3 (P/N 1000078, 10x Genomics). Sequencing libraries were loaded at 2.4 pM on Illumina NovaSeq 6000 with 2×150 paired-end kits using the following read length: 28 bp Read1, 8 bp I7 Index, 8 bp I5 Index and 91 bp Read2.

### scRNA-seq data of bead-enriched T cell subsets

We downloaded scRNA-seq data of 6 bead-enriched T cell subsets and PBMCs from the website of 10X GENOMICS (https://support.10xgenomics.com/single-cell-gene-expression/datasets). The 6 bead-enriched T cell subsets are CD4+ T helper Cells (CD4 Th), CD4+/CD45RA+/CD25− naive T cells (CD4 Naive), CD4+/CD25+ Regulatory T Cells (CD4 Treg), CD4+/CD45RO+ Memory T Cells (CD4 Memory), CD8+ cytotoxic T cells (CD8 CTL), and CD8+/CD45RA+ naive T cells (CD8 Naive). FACS analyses showed the high purity of those these T cell subsets: CD4 T (99%), CD4 Naïve (98%), CD4 Memory (98%), CD4 Treg (95%), CD8 Naïve (99%) and CD8 CTL (98%). These T cell subsets were processed by 10X Genomics Chromium with the aim of capturing 10,000 cells for each sample. The basic sample information also is available in Zheng *et al.*(*22*).

### Data pre-processing and quality control (QC)

Cell ranger was used for generating the single cell expression matrix. We filtered out the cells meeting any of the following criterion: 1) less than 200 unique genes, 2) more than 1,500 unique genes, 3) >50,00 UMI counts, 4) more than 5% mitochondrial counts. After QC, we normalized expression level of each gene to the total number of UMI for each cell.

### Dimension reduction and visualization of scRNA-seq data

Seurat (*23*) is used for data integration, data normalization, dimension reduction, cell clustering and other basic scRNA-seq data analyses following our previous studies (*5, 15*). We calculated the cell-to-cell variation of each gene and selected the top 2,000 genes exhibiting the highest variation for further analysis. To avoid highly expressed genes dominate in later analyses, we scaled the mean and variance of each gene across cells is 0 and 1, respectively. t-distributed stochastic neighbor embedding (t-SNE) (*15, 37, 38*) in Seurat was used for data visualization. We displayed cluster specific expression genes on t-SNE, which provide nice visualization for distinguishing different cell clusters.

### Gene Ontology (GO) enrichment analysis

To examine whether particular GO terms were enriched in certain gene set, we carried out GO enrichment analysis using DAVID (*39*). GO categories with FDR < 0.05 were consider as significant.

### Relationships between bead-enriched T cell subsets and scCPops

Sankey diagram is an intuitive visualization approach for depicting flows from one set of values to another. We plotted Sankey diagram using Highcharts (https://www.highcharts.com/demo/sankey-diagram) to explore the relationships between bead-enriched T cell subsets and scCPops.

### Single cell trajectories inference

We implemented Slingshot (*29*) on scRNA-seq data to infer the trajectory of CD4 T cells and CD8 T cells. The scCPops were used as input for Slingshot to infer the pseudotime. Cell lineages were visualized on the PCA three-dimensional diagram using plot3D in R.

### Statistical analysis and key Reagents

The statistical tests and plots were conducted using R version 3.5.2 (2018-12-20) (*40*). The key reagent resources for this study are available (Table S3).

## Data availability

The raw sequence data reported in this paper have been deposited in the Genome Sequence Archive in BIG Data Center, under accession numbers HRA000313.

## Acknowledgements

We thank Dr. Keji Zhao for critical reading of the manuscript.

## Funding

This study was supported by National Key R&D Program of China (2018YFC1004500), National Natural Science Foundation of China (81872330), the Science and Technology Innovation Commission of the Shenzhen Municipal Government (JCYJ20170817111841427), the Shenzhen Science and Technology Program (KQTD20180411143432337), and Center for Computational Science and Engineering, Southern University of Science and Technology. The funders had no role in study design, data collection and analysis, decision to publish, or preparation of the manuscript.

## Author’s contributions

W.J. conceived and designed the project. S.C. did the experiment. X.W. and X.S. analyzed the data, with contribution from H.L.. W.J., N.H. and X.C. supervise the study. W.J. and X. W. wrote the manuscript. All authors have read and approved the manuscript.

## Conflict of Interests

The authors declare no conflict of interests.

## References

1. L. Zhang et al., Lineage tracking reveals dynamic relationships of T cells in colorectal cancer. Nature 564, 268–272 (2018).

2. K. K. Farh et al., Genetic and epigenetic fine mapping of causal autoimmune disease variants. Nature 518, 337–343 (2015).

3. D. L. Farber, N. A. Yudanin, N. P. Restifo, Human memory T cells: generation, compartmentalization and homeostasis. Nat Rev Immunol 14, 24–35 (2014).

4. G. Hu et al., Transformation of Accessible Chromatin and 3D Nucleome Underlies Lineage Commitment of Early T Cells. Immunity 48, 227–242 e228 (2018).

5. P. Qin et al., Integrated decoding hematopoiesis and leukemogenesis using single-cell sequencing and its medical implication. Cell Discov 7, 2 (2021).

6. J. Zhu, W. E. Paul, CD4 T cells: fates, functions, and faults. Blood 112, 1557–1569 (2008).

7. T. R. Mosmann, H. Cherwinski, M. W. Bond, M. A. Giedlin, R. L. Coffman, Two types of murine helper T cell clone. I. Definition according to profiles of lymphokine activities and secreted proteins. J. Immunol. 136, 2348–2357 (1986).

8. S. Sakaguchi, N. Sakaguchi, M. Asano, M. Itoh, M. Toda, Immunologic self-tolerance maintained by activated T cells expressing IL-2 receptor alpha-chains (CD25). Breakdown of a single mechanism of self-tolerance causes various autoimmune diseases. J Immunol 155, 1151–1164 (1995).

9. J. D. Fontenot, M. A. Gavin, A. Y. Rudensky, Foxp3 programs the development and function of CD4(+)CD25(+) regulatory T cells. Nat. Immunol. 4, 330–336 (2003).

10. M. A. Suni et al., CD4(+)CD8(dim) T lymphocytes exhibit enhanced cytokine expression, proliferation and cytotoxic activity in response to HCMV and HIV-1 antigens. Eur. J. Immunol. 31, 2512–2520 (2001).

11. A. M. Intlekofer et al., Effector and memory CD8(+) T cell fate coupled by T-bet and eomesodermin. Nat. Immunol. 6, 1236–1244 (2005).

12. W. Wang, G. Ren, N. Hong, W. Jin, Exploring the changing landscape of cell-to-cell variation after CTCF knockdown via single cell RNA-seq. BMC Genomics 20, 1015 (2019).

13. E. Z. Macosko et al., Highly Parallel Genome-wide Expression Profiling of Individual Cells Using Nanoliter Droplets. Cell 161, 1202–1214 (2015).

14. F. Paul et al., Transcriptional Heterogeneity and Lineage Commitment in Myeloid Progenitors. Cell 163, 1663–1677 (2015).

15. B. Zhou, W. Jin, Visualization of Single Cell RNA-Seq Data Using t-SNE in R. Methods Mol Biol 2117, 159–167 (2020).

16. Y. Yu et al., Single-cell RNA-seq identifies a PD-1(hi) ILC progenitor and defines its development pathway. Nature 539, 102–106 (2016).

17. V. Proserpio et al., Single-cell analysis of CD4+ T-cell differentiation reveals three major cell states and progressive acceleration of proliferation. Genome Biol 17, 103 (2016).

18. A. T. Waickman et al., Dissecting the heterogeneity of DENV vaccine-elicited cellular immunity using single-cell RNA sequencing and metabolic profiling. Nat Commun 10, 3666 (2019).

19. D. Zemmour et al., Single-cell gene expression reveals a landscape of regulatory T cell phenotypes shaped by the TCR. Nat Immunol 19, 291–301 (2018).

20. V. S. Patil et al., Precursors of human CD4(+) cytotoxic T lymphocytes identified by single-cell transcriptome analysis. Sci Immunol 3, (2018).

21. P. K. Chattopadhyay, T. M. Gierahn, M. Roederer, J. C. Love, Single-cell technologies for monitoring immune systems. Nat Immunol 15, 128–135 (2014).

22. G. X. Zheng et al., Massively parallel digital transcriptional profiling of single cells. Nat Commun 8, 14049 (2017).

23. A. Butler, P. Hoffman, P. Smibert, E. Papalexi, R. Satija, Integrating single-cell transcriptomic data across different conditions, technologies, and species. Nat Biotechnol 36, 411–420 (2018).

24. T. Stuart et al., Comprehensive Integration of Single-Cell Data. Cell 177, 1888–1902 e1821 (2019).

25. O. Van Grembergen et al., Portraying breast cancers with long noncoding RNAs. Sci Adv 2, e1600220 (2016).

26. J. Zhou et al., Linc00152 promotes proliferation in gastric cancer through the EGFR-dependent pathway. J Exp Clin Cancer Res 34, 135 (2015).

27. T. Hayakawa, K. Yamashita, K. Tanzawa, E. Uchijima, K. Iwata, Growth-promoting activity of tissue inhibitor of metalloproteinases-1 (TIMP-1) for a wide range of cells. A possible new growth factor in serum. FEBS Lett 298, 29–32 (1992).

28. P. R. Rogers, J. Song, I. Gramaglia, N. Killeen, M. Croft, OX40 promotes Bcl-xL and Bcl-2 expression and is essential for long-term survival of CD4 T cells. Immunity 15, 445–455 (2001).

29. K. Street et al., Slingshot: cell lineage and pseudotime inference for single-cell transcriptomics. BMC Genomics 19, 477 (2018).

30. K. Hashimoto et al., Single-cell transcriptomics reveals expansion of cytotoxic CD4 T cells in supercentenarians. Proc Natl Acad Sci U S A 116, 24242–24251 (2019).

31. E. Cano-Gamez et al., Single-cell transcriptomics identifies an effectorness gradient shaping the response of CD4(+) T cells to cytokines. Nat Commun 11, 1801 (2020).

32. S. Urata et al., BST-2 controls T cell proliferation and exhaustion by shaping the early distribution of a persistent viral infection. PLoS Pathog 14, e1007172 (2018).

33. B. Oriol-Tordera et al., Methylation regulation of Antiviral host factors, Interferon Stimulated Genes (ISGs) and T-cell responses associated with natural HIV control. PLoS Pathog 16, e1008678 (2020).

34. K. Kisand et al., Interferon autoantibodies associated with AIRE deficiency decrease the expression of IFN-stimulated genes. Blood 112, 2657–2666 (2008).

35. S. C. Jameson, D. Masopust, Understanding Subset Diversity in T Cell Memory. Immunity 48, 214–226 (2018).

36. S. Hoyer et al., Concurrent interaction of DCs with CD4(+) and CD8(+) T cells improves secondary CTL expansion: It takes three to tango. Eur J Immunol 44, 3543–3559 (2014).

37. L. van der Maaten, G. Hinton, Visualizing Data using t-SNE. J Mach Learn Res 9, 2579–2605 (2008).

38. A. Mahfouz et al., Visualizing the spatial gene expression organization in the brain through non-linear similarity embeddings. Methods 73, 79–89 (2015).

39. W. Huang da, B. T. Sherman, R. A. Lempicki, Systematic and integrative analysis of large gene lists using DAVID bioinformatics resources. Nature protocols 4, 44–57 (2009).

40. R Core Team, R: A Language and Environment for Statistical Computing. (R Foundation for Statistical Computing, Vienna, Austria, 2018).

